# A Resource of Gene Expression Data from a Multiethnic Population Cohort of Induced Pluripotent Cell–Derived Cardiomyocytes

**DOI:** 10.1101/2023.05.15.539274

**Authors:** Wenjian Lv, Apoorva Babu, Michael P. Morley, Kirin Musunuru, Marie Guerraty

## Abstract

Induced pluripotent stem cells (iPSCs), upon differentiation into somatic cell types, offer the ability to model the genetic states of tissues in the individuals from whom the iPSCs were derived.^1^ A prior study used a multiethnic cohort of healthy individuals to derive iPSCs and iPSC-hepatocytes and perform gene expression and expression quantitative trait locus (eQTL) analyses, identifying causal genes and variants linked to blood lipid levels.^2^ We sought to use this cohort of iPSC lines to generate a similar resource with iPSC-cardiomyocytes (iPSC-CMs) and make the results available to the community.

## METHODS

As previously described, mononuclear cells isolated from peripheral blood samples of 91 healthy individuals were used to generate iPSC lines.^2^ We differentiated the various cell lines (one per individual) into iPSC-CMs using an established 2-dimensional differentiation protocol.^3,4^ In brief, we used feeder-free differentiation conditions entailing the addition of a variety of growth factors and chemicals to the growth media. Undifferentiated iPSCs were detached by a 4-min incubation with Accutase (StemCell technologies) and seeded onto Geltrex (Life Technologies)-coated plates at a density of 200,000 cells/cm^2^. To induce cardiac differentiation, we replaced iPSC Brew (StemMACS), in which the iPSCs had been maintained, with RPMI-B27 [RPMI-1640 (Life Technologies); 2% B-27 Supplement Minus Insulin (B27-Insulin) (Thermo Fisher Scientific)] medium supplemented with recombinant human/mouse/rat activin A (100 ng/mL; R&D Systems) for 18 hours, followed by recombinant human BMP-4 (5 ng/mL; R&D Systems) and CHIR 99021 (1 nM, Tocris) for 2 days. The medium was then exchanged for RPMI-B27-Insulin with Xav (1nM, Tocris) for another 2 days followed by RPMI-B27-Insulin without supplementary cytokines for 3 more days. RPMI/B-27-Insulin was refed every 2–3 days for several additional weeks. Widespread spontaneous beating activity was typically observed by day 10 after activin A addition. They then underwent 2 3-day rounds of metabolic selection in glucose-free media, followed by recovery, to eliminate non-cardiomyocytes. The cells were maintained until day 30, when they were analyzed by flow cytometry and harvested for RNA for sequencing.

## RESULTS

We found that 71 out of 91 iPSC lines successfully differentiated into iPSC-CM lines (34 from individuals self-identified as African-Americans and 37 from individuals self-identified as European-Americans; 44 from women, 32 from men), defined as ≥80% positive for troponin T staining (AB-8295, abcam) using flow cytometry (Figure, A). Total RNA was isolated using the RNeasy Mini Kit (QIAGEN) followed by oligo-dT selection. RNA sequencing libraries were prepared using the Illumina TruSeq stranded mRNA kit followed by the Nugen Ovation amplification kit. Libraries were sequenced in 3 batches with an Illumina HiSeq2500 system (performed on a fee-for-service basis by GENEWIZ, Inc.) to a depth of ≈30 million 150-bp paired-end reads per biological sample. FASTQ files were aligned against human reference (hg19/hGRC37) using the STAR aligner. Duplicate reads were removed using MarkDuplicates from Picard tools, and per gene read counts for Ensembl (v75) gene annotations were computed. Expression levels in counts per million (CPM) were normalized and transformed using VOOM in the LIMMA R package. To account for observed batch effects, a surrogate variant analysis was performed using the R package SVAseq and 11 additional covariates were identified. LIMMA was used perform differential gene expression analysis. Expression quantitative trait locus analysis was performed using the QTLtools package with adjustment for sex, race, and the first 3 genetic principal components and the 11 SVA-computed covariates.

**Figure.**
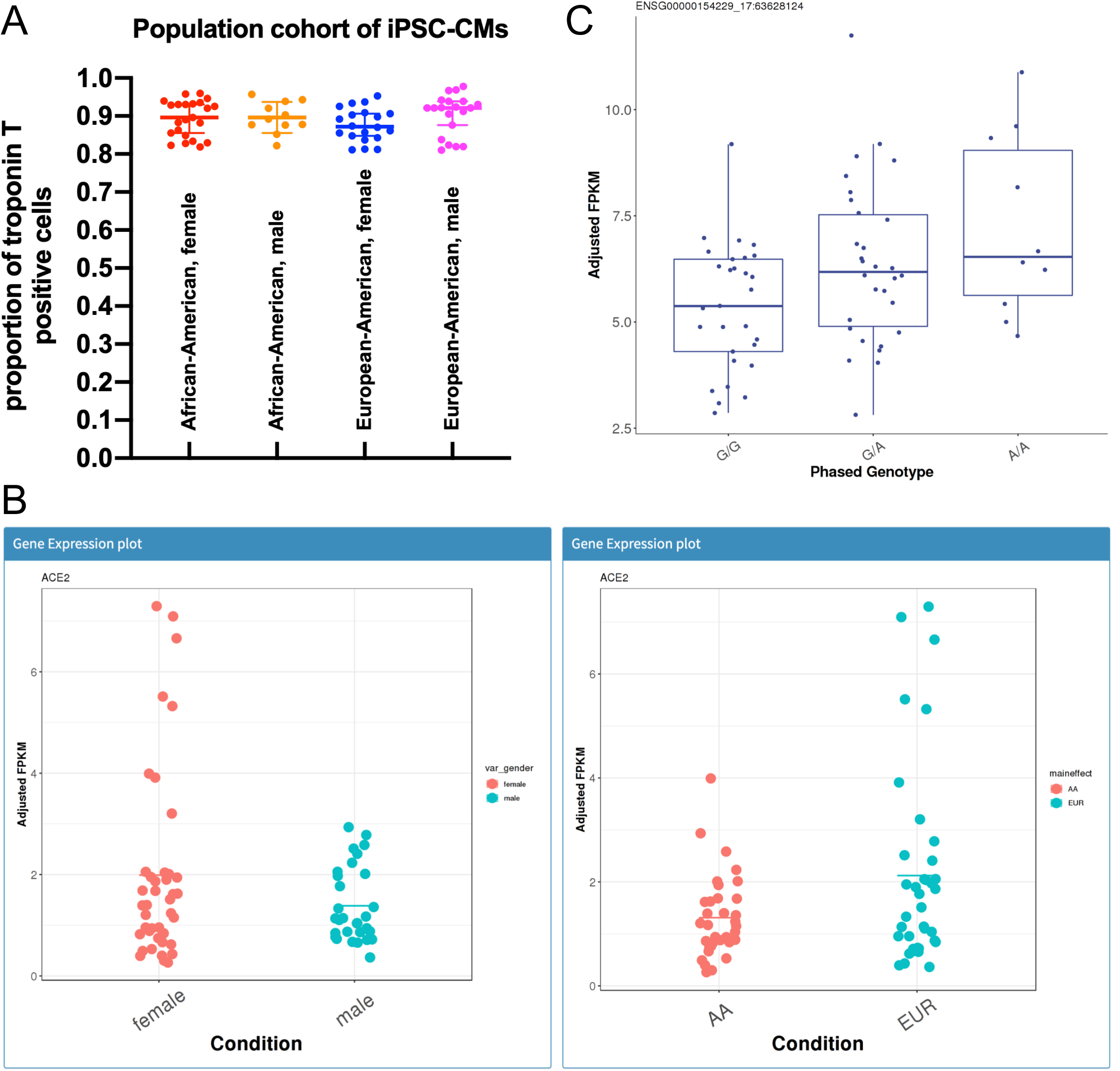
A resource of gene expression data from a multiethnic population cohort of iPSC-CMs. **A**, Troponin T positivity of iPSC-CMs as a marker of differentiation, with iPSC-CM samples stratified by gender and race. Bars indicate means and standard deviations. **B**, Sample snapshots from the resource website documenting differences in *ACE2* expression by gender and race. **C**, Sample snapshot displaying an eQTL relationship between *PKCRA* and rs149582052.

The iPSC-CM gene expression and eQTL data resource is freely accessible via the website http://165.123.69.6:3838/ipsc/. A catalog of the iPSC-CM lines provides self-identified gender, race, and age and is otherwise de-identified. The searchable gene expression database has the ability to display expression levels of individual genes as dot blots, offering comparisons by gender (male versus female) and race (European-American versus African-American). For example, *ACE2* expression in iPSC-CMs varies by both gender and ancestry (Figure, B), highlighting the importance of considering these variables in the interpretation of iPSC-CM data. The same comparisons can be applied across all genes and displayed as volcano plots or downloaded as tables with effect sizes and *P*-values. All of the raw expression data can be downloaded in tabular format.

The eQTL database can be queried for genotype-expression data with an individual gene name or an individual DNA variant (rs number), generating a dot plot against the DNA variant or the gene, respectively, with the strongest statistically significant eQTL association in the aggregate iPSC-CM samples. See Figure, C, for a snapshot from the resource website for the established left ventricular eQTL gene *PRKCA* (protein kinase C-α). There is also the ability to generate a dot plot for any desired combination of gene and DNA variant, assuming both are present in the database.

## CONCLUSIONS

We anticipate this multiethnic resource of iPSC-CM gene expression and eQTL data will be broadly useful to the cardiovascular research community, both in facilitating functional genomics studies and in assessing for gender and interethnic differences in cardiomyocyte expression levels of genes of physiological interest.^5^

## Data Sharing

The RNA-seq data and genotype data used to generate this resource are available via the accession numbers GSE146071 and dbGaP: phs001341.v1.p1, respectively.

## Sources of Funding

This work was supported by funds from the University of Pennsylvania, the Winkelman Family Fund in Cardiovascular Innovation (WL), NHLBI (MG) and Burroughs Wellcome Fund (MG).

## Disclosures

None.

## Notes

http://165.123.69.6:3838/ipsc/

### Competing Interest Statement

The authors have declared no competing interest.

